# AB-Free Kava Suppresses Tobacco Smoke-Induced CREB Phosphorylation in the Mouse Cerebellum

**DOI:** 10.1101/2025.03.18.643897

**Authors:** Yifan Wang, Tara Hashemian, Tengfei Bian, Leah R. Reznikov, Adriaan W. Bruijnzeel, Tuo Lin, Chengguo Xing

## Abstract

**Introduction:** Our recent studies revealed the potential of AB-free kava as a novel therapeutic candidate against tobacco use disorder while the underlying mechanism remains to be elucidated. The cerebellum region in the brain has recently been reported to be involved in tobacco smoke withdrawal. The cAMP response element-binding protein (CREB) activation via phosphorylation has also been reported to contribute to substance abuse, including tobacco use disorder. Our early pre-clinical work revealed the potential of AB-free kava to suppress tobacco smoke-induced CREB activation in lung tissues.

**Methods:** In this study, we investigated the impact of tobacco smoke exposure on the levels of phosphorylated CREB (p-CREB) in the mouse cerebellum and evaluated the effects of AB-free kava.

**Results:** Four different regimens of tobacco smoke exposure consistently increased the levels of p-CREB in the cerebellum. Importantly, AB-free kava effectively suppressed tobacco smoke-induced increase in p-CREB to levels comparable to mice without tobacco smoke exposure.

**Conclusions:** CREB activation in the cerebellum is a potential mechanism involved in tobacco use disorder, and our data provide preliminary evidence for the protective potential of AB-free kava.

## Introduction

Tobacco smoke is one of the leading causes of morbidity and mortality worldwide,^1^ contributing to a range of preventable diseases that extend far beyond the respiratory system.^2^ For instance, prolonged tobacco smoke has been shown to perturb brain structure and regulation, leading to deleterious outcomes, including nicotine addiction, cognitive impairments, and elevated risks of stroke.^3^ Understanding the underlying molecular mechanisms and finding effective interventions remain a critical area of research.

Kava (*Piper methysticum*) is a plant native to the South Pacific Islands. Its root can be processed into a beverage or a dietary supplement that has been used to reduce stress and promote relaxation.^4^ Certain kava formulations, particularly those prepared with low-quality raw materials not recommended for human consumption through organic extraction, contain high levels of flavokavains A and B, which may pose an increased hepatotoxic risk.^5^ To address this potential limitation, an AB-free kava formula was developed, composed mainly of six major kavalactones with flavokavains A and B removed.^6,7^ Our prior preclinical research demonstrated the potential of AB-free kava in mitigating several tobacco smoke-related health issues.^7,8^ For instance, AB-free kava and one of its kavalactones (dihydromethysticin) at a human exposure relevant dose effectively prevented tobacco-specific carcinogen-induced lung carcinogenesis in A/J mice partly by suppressing the protein kinase A (PKA) mediated cAMP response element-binding protein (CREB) signaling pathway in the lung.^7,9^ Additionally, dietary administration of AB-free kava revealed a unique global transcriptional neutralization in the lung and liver in tobacco smoke-exposed mice.^8^ Part of its global protective potential against tobacco smoke was highlighted in somatic withdrawal symptoms.^8^ Consistently, our results revealed its potential to reduce tobacco use and dependence even among addicted smokers with no intention to quit.^10^ More importantly, CREB phosphorylation has also been reported to play a central role in anxiety regulation^11^ and nicotine addiction.^12^ This connection further aligns with the potential benefit of using kava to manage anxiety and tobacco use.

The cerebellum, while best characterized for its role in motor coordination,^13^ has been increasingly recognized for its potential contributions to addiction.^14,15^ Mechanistically, nicotine exerts its effects not only by activating nicotinic acetylcholine receptors but also by elevating adenylyl cyclase activity, a key enzyme activating PKA/CREB signaling, in the cerebellum.^16^ These findings highlight the underexplored vulnerability of cerebellum to tobacco use disorder and the potential involvement of the PKA/CREB signaling.

Given the well-known functions of kava in anxiety management, its potential to reduce tobacco dependence,^8,10^ and its effects on CREB in the lung tissues, this study characterized the effects of tobacco smoke on CREB phosphorylation in mouse cerebellum and the counteracting potential of AB-free kava, which may provide mechanistic insight for kava’s potential against tobacco use disorder.

## Materials and Methods

### Caution

NNK and tobacco smoke are Class IA human carcinogens and should be handled carefully in well-ventilated fume hoods with proper protective clothing.

#### Chemicals and Reagents

The AIN-93M powdered diets were purchased from Harlan Teklad (Cambridgeshire, UK). Diet supplemented with AB-free kava at the specified dose was prepared, characterized, stored and dispensed as previously described.^7^ 1R6F research cigarettes were purchased from Tobacco and Health Research Institute, University of Kentucky. SDS-PAGE gel was purchased from GenScript (#M00654). The antibodies were Phospho-CREB (Ser133) (87G3) Rabbit mAb (Cell Signaling Technology, #9198), Anti-rabbit IgG, HRP-linked Antibody (Cell Signaling Technology, #7074), and Mouse monoclonal anti-β-actin (Sigma Aldrich, #A2228).

#### Animal Studies

Female A/J mice were purchased from the Jackson Laboratory (Bar Harbor, ME) and maintained in the specific pathogen-free facilities, according to animal welfare protocols approved by Institutional Animal Care and Use Committee at the University of Florida. Mice, randomized into different groups based on body weights after 7 days of acclimation, were exposed to filtered air or tobacco smoke as described below. Mice were maintained on AIN-M powdered diet with or without AB-free kava supplement at a dose of 0.75 mg/g or 0.25 mg/g of diet as previously reported.^8^ After the experiment cycle, mice were euthanized with the cerebellum collected and stored at -80°C till analysis. Four rounds of mouse studies were conducted with different tobacco smoke exposure regimens, which were designed to mimic different tobacco smoke exposures in human smokers (Fig. 1A).

**Figure. 1.**
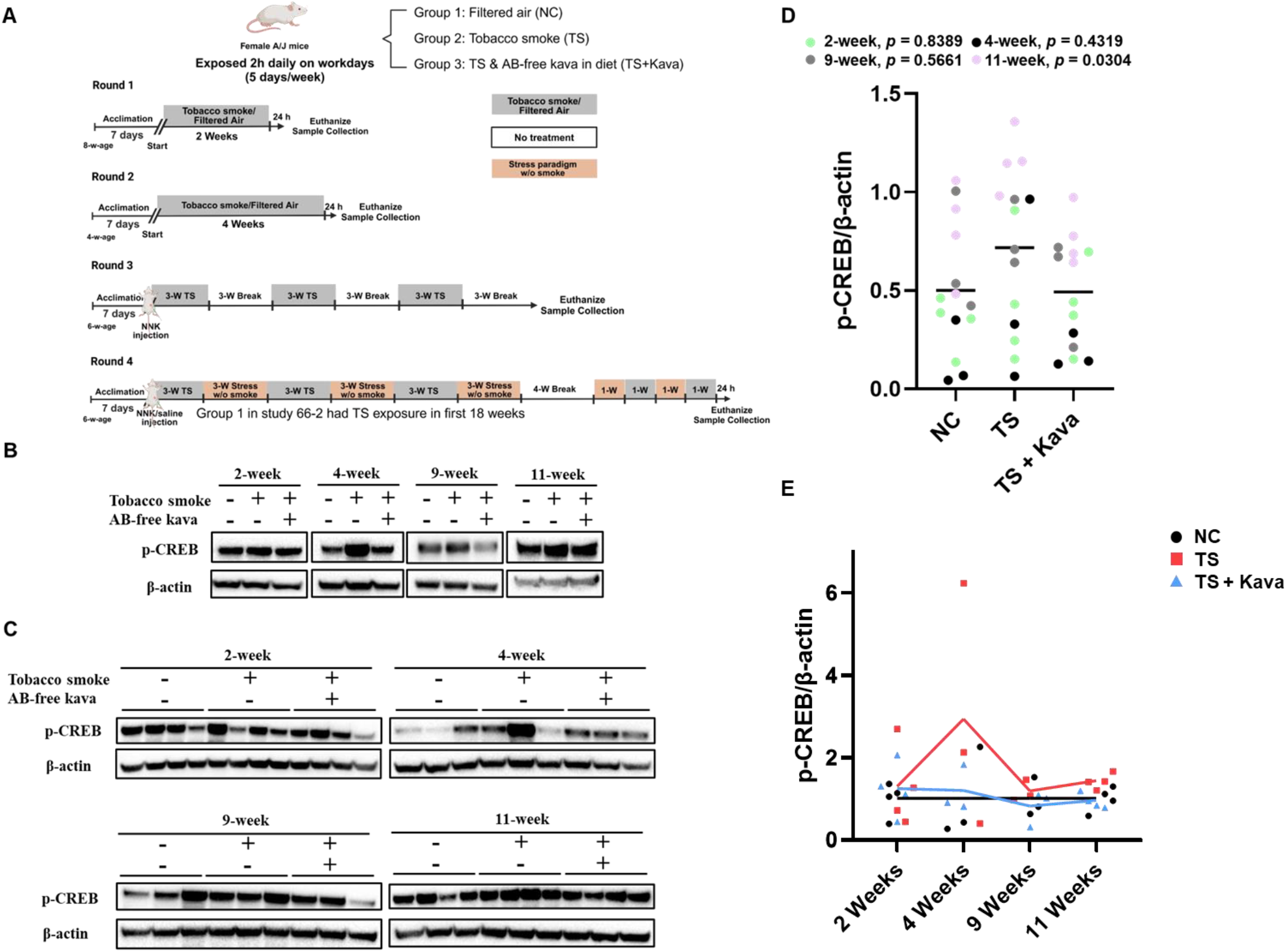
TS induced elevated CREB phosphorylation in the cerebellum with AB-free kava effective suppression. (A) Tobacco smoke exposure animal studies scheme. Western blot analysis of cerebellum p-CREB upon TS exposure with/without AB-free kava supplementation, shown as (B) pooled samples, (C) individual samples, and (D) quantitative visualization. (E) p-CREB/β-actin levels changes over time with respect to exposure (NC – air exposure; TS – tobacco smoke, TS + Kava – tobacco smoke + AB-free kava).

#### Tobacco Smoke Exposure

Tobacco smoke exposure sessions were carried out as previously described.^17^Mice were exposed to tobacco smoke using a microprocessor-controlled smoking machine (TE-10, Teague Enterprises) under a standardized procedure (35 cm^3^ puff volume, 1 puff/min, 2 s per puff, 8 puffs/cigarette) for 2 h daily (9–11 am, Monday to Friday), with total suspended particles (TSP) at ∼100 mg/m^3^. TSP and CO levels were measured using gravimetric sampling and a CO analyzer (Monoxor III).

#### Stress Paradigm

To mimic human smokers who experience different forms and levels of stress, mice were subjected to a chronic unpredictable stress paradigm following reported procedures.^18^ Mice were moved to a designated room and subjected to one stressor from paradigm list (wet cage, dampened bedding, tilted cage, etc., Supplementary materials) at a random time 3 times/week (Monday, Wednesday, Friday).

#### Western Blotting

Around 30 mg of cerebellum tissue was homogenized in 500 µL RIPA buffer. After centrifugation (13,000 x g, 15 min, 4°C), the supernatant was collected with protein concentration determined by BCA assay. Then, 50 µg protein was separated on a 4-12% SDS-PAGE gel, transferred to a PVDF membrane, blocked with 5% milk, and incubated overnight at 4°C with primary antibody, followed by an HRP-conjugated secondary antibody. Proteins were detected using ECL reagent and imaged using a Bio-Rad system.

#### Image Quantitative Analysis

Western blot images were quantified using ImageJ. Blot images were converted to 8-bit; background subtraction (rolling ball radius: 50 pixels) was applied; and bands were selected using the rectangular tool. Area and integrated density were measured, with the scale set to 1.0 pixel. Densitometric analysis (‘Analyze → Gels → Plot Lanes’) generated intensity profiles, and band intensities were normalized to β-actin for statistical analysis of relative protein levels.

### Statistical Analysis

Given that p-CREB/β-actin values were collected from four rounds of animal studies, the linear mixed-effect model (LMM) was used to assess the association between the p-CREB protein levels and interventions. The LMM included a fixed intervention effect and a random intercept effect to address inter-cluster variability, followed by post-hoc pairwise comparisons using the false discovery rate (FDR) method to adjust for multiple comparisons.

## Results and Discussions

With varied durations of tobacco smoke exposure, a consistent increase in p-CREB protein level was observed in mouse cerebellum in comparison to mice with air exposure (Fig. 1B). The levels of total CREB remained the same (representative data in Fig. S1). The increase in p-CREB levels was noticed as early as two weeks of tobacco smoke exposure when somatic withdrawal symptoms started to develop.^8^ The consistent p-CREB increases observed with different smoke exposure regimens strongly support the effect of tobacco smoke on CREB activation in mouse cerebellum. Notably, the increased p-CREB levels was effectively suppressed by AB-free kava dietary supplement under all conditions evaluated, supporting the neutralizing potential of AB-free kava against tobacco smoke. To assess the extent of individual variations, p-CREB levels were analyzed in individual mouse cerebellum samples (Fig. 1C). While high levels of heterogeneity were observed between individual mice, the overall trend of increased p-CREB levels in tobacco smoke-exposed groups and the suppressive effects of AB-free kava were consistent across the samples. Quantitative analyses in each round were mostly not statistically significant by using One-way ANOVA, due to the limited sample size in each group (n=3 or 4) and the high individual mouse heterogeneity (Fig. 1D). To address such limitations, we implemented a linear mixed-effect model (LMM) to analyze the four rounds of mouse studies collectively (Table 1). The LMM combines fixed and random effects that account for within-cluster/batch correlations. The results revealed that tobacco smoke markedly increased p-CREB levels in the cerebellum (p = 0.03). Furthermore, the increased p-CREB levels were effectively suppressed by AB-free kava (p = 0.03), demonstrating its potential protective effects.

**Table. 1.** Pairwise comparisons of p-CREB/β-actin levels value using LMM.

| **Contrast** | **Estimate** | **95% CI** | **p-value** |
| --- | --- | --- | --- |
| TS - NC | 0.22 | 0.05, 0.39 | **0.03** |
| TS - (TS+Kava) | 0.23 | 0.06, 0.40 | **0.03** |
| NC - (TS+Kava) | 0.01 | -0.16, 0.18 | 0.93 |

NC – air exposure; TS – tobacco smoke; TS + Kava – tobacco smoke + AB-free kava; 95% CI – 95% confidence interval

To further explore the dynamics of p-CREB changes in response to tobacco smoke and the potential mitigating effects of AB-free kava, we investigated p-CREB levels across different tobacco exposure durations. To account for variability between experimental batches, the p-CREB levels from all three groups (NC, TS, TS+Kava) were normalized to the mean value of the negative control (NC) group within each experimental batch. The data revealed a distinct pattern of p-CREB changes over time (Fig. 1E). Specifically, increased p-CREB levels were observed with prolonged tobacco smoke exposure (Week 2 and 4), suggesting a time-dependent effect. In the 9-week study, however, this increasing trend reversed, with p-CREB levels decreasing. This likely reflects the cessation of tobacco smoke exposure, as cerebellum samples were collected three weeks after the final exposure. Notably, when tobacco smoke was reintroduced following the cessation period, p-CREB levels increased again (Week 11), indicating that p-CREB changes in the cerebellum may be reversible but can be reactivated upon tobacco smoke re-exposure. The reversibility of p-CREB provides insight into the dynamic nature of p-CREB regulation in the cerebellum.^19^ An important and consistent observation was that AB-free kava mitigated the tobacco smoke-induced increases in p-CREB levels across all four regimens, which again supports its potential as a protective agent.

As discussed, p-CREB has been implicated in anxiety and addiction.^11,12^ Emerging evidence also highlights cerebellum’s role in addiction.^15^ Tobacco smoke-induced p-CREB alterations in the cerebellum may contribute to the development and maintenance of tobacco use disorder. AB-free kava may reduce tobacco dependence by suppressing tobacco smoke-induced p-CREB in the cerebellum. In addition, mice in these four rounds have different smoking exposures and other variations, such as exposure to the additional carcinogen (NNK) and stress, at least partially mimicking different life experiences among human smokers. With the same trend of p-CREB changes observed, our data strongly supports the significant effects of tobacco smoke on p-CREB in the cerebellum and the suppressive effects of AB-free kava.

This study has several limitations. First, while assessing p-CREB protein provides molecular insights, these studies did not evaluate potential behavioral changes. Thus, the functional impact from p-CREB remains to be determined. Second, p-CREB may be involved in other brain regions for tobacco use disorder and other neurological functions, which requires further characterization. Lastly, the studies were only performed in female mice of similar age in a single strain. Its scope remains to be characterized.

## Conclusion

The cerebellum is an underexplored region for tobacco use disorder. This study demonstrated that tobacco smoke exposure leads to a significant increase in p-CREB levels in the mouse cerebellum, which may contribute to tobacco use disorder. Notably, AB-free kava dietary supplements effectively and consistently suppressed tobacco smoke-induced p-CREB elevation. These findings support the potential of AB-free kava in mitigating tobacco smoke-induced effects in the brain, such as tobacco use disorder.

## Supplementary Materials

Supplementary material is available at *Nicotine and Tobacco Research* online.

## Funding

This research was supported by a grant from the Florida Department of Health (23B02, CX).

## Declaration of Interests

The authors declare the following competing financial interest(s): the AB-free kava product evaluated is based on the IPs owned by Kuality Herbceutics (KH). There is no profit or product from KH yet. Chengguo Xing has a 55% ownership of KH. He also provides consultation to KH for its strategy on development. All other authors declare no conflict of interest.

### Acknowledgment

We appreciate Dr. Jessica Mamallapalli, Breanne Freeman, Allison Lynch, Kayleigh Ballas, and Alexander Scala contributing to the animal tobacco smoke exposure studies, and technical support from Dr. Brandon Warren.

## Author Contributions

Yifan Wang (Conceptualization [lead], Formal analysis [lead], Writing – original draft [lead], Writing – review & editing [lead]), Tara Hashemian (Formal analysis [lead], Writing – original draft [equal], Writing – review & editing [equal]), Tengfei Bian (Formal analysis [supporting], Writing – review & editing [equal]), Leah R. Reznikov(Formal analysis [supporting], Writing – review & editing [equal]), Adriaan W. Bruijnzeel (Formal analysis [supporting], Resources [lead], Writing – review & editing [equal]), Tuo Lin (Formal analysis [lead], Writing – review & editing [equal]), Chengguo Xing (Funding acquisition [lead], Conceptualization [lead], Resources [lead], Supervision [lead], Writing – original draft [lead], Writing – review & editing [lead])

## Data Availability

The data of this manuscript are available from the corresponding author.

## References

1. Yang JJ, Yu D, Wen W, et al. Tobacco Smoking and Mortality in Asia: A Pooled Meta-analysis. JAMA Netw Open. 2019;2(3):e191474.

2. Carter BD, Abnet CC, Feskanich D, et al. Smoking and mortality--beyond established causes. N Engl J Med. 2015;372(7):631–631.

3. Le Foll B, Piper ME, Fowler CD, et al. Tobacco and nicotine use. Nat Rev Dis Primers. 2022;8(1):19.

4. Bian T, Corral P, Wang Y, et al. Kava as a Clinical Nutrient: Promises and Challenges. Nutrients. 2020;12(10).

5. Narayanapillai SC, Leitzman P, O’Sullivan MG, Xing C. Flavokawains a and B in kava, not dihydromethysticin, potentiate acetaminophen-induced hepatotoxicity in C57BL/6 mice. Chem Res Toxicol. 2014;27(10):1871–1871.

6. Narayanapillai SC, Balbo S, Leitzman P, et al. Dihydromethysticin from kava blocks tobacco carcinogen 4-(methylnitrosamino)-1-(3-pyridyl)-1-butanone-induced lung tumorigenesis and differentially reduces DNA damage in A/J mice. Carcinogenesis. 2014;35(10):2365–2365.

7. Leitzman P, Narayanapillai SC, Balbo S, et al. Kava blocks 4-(methylnitrosamino)-1-(3-pyridyl)-1-butanone-induced lung tumorigenesis in association with reducing O6-methylguanine DNA adduct in A/J mice. Cancer Prev Res. 2014;7(1):86–86.

8. Bian T, Lynch A, Ballas K, et al. Flavokavains A- and B-Free Kava Enhances Resilience against the Adverse Health Effects of Tobacco Smoke in Mice. ACS Pharmacol Transl Sci. 2024;7(11):3502–3517.

9. Bian T, Ding H, Wang Y, et al. Suppressing the activation of protein kinase A as a DNA damage-independent mechanistic lead for dihydromethysticin prophylaxis of NNK-induced lung carcinogenesis. Carcinogenesis. 2022;43(7):659–659.

10. Wang YN, S.,,,,,, Tessier, K.,,,,,, Strayer, L.,,,,,, Upadhyaya, P.,,,,,, Hu, Q.,,,,,, Kingston, R.,,,,,, Salloum, R. G.,,,,,, Lu, J.,,,,,, Hecht, S.S.,,,,,, Hatsukami, D.,,,,,, Fujioka, N.,,,,,, Xing, C. The Impact of One-week Dietary Supplementation with Kava on Biomarkers of Tobacco Use and Nitrosamine-based Carcinogenesis Risk among Active Smokers. Cancer Prev Res. 2020;13(5):483–483.

11. Zhang J, Cai CY, Wu HY, Zhu LJ, Luo CX, Zhu DY. CREB-mediated synaptogenesis and neurogenesis is crucial for the role of 5-HT1a receptors in modulating anxiety behaviors. Sci Rep. 2016;6:29551.

12. Fisher ML, LeMalefant RM, Zhou L, Huang G, Turner JR. Distinct Roles of CREB Within the Ventral and Dorsal Hippocampus in Mediating Nicotine Withdrawal Phenotypes. Neuropsychopharmacology. 2017;42(8):1599–1599.

13. Ito M. Mechanisms of motor learning in the cerebellum. Brain Res. 2000;886(1-2):237–245.

14. Moulton EA, Elman I, Becerra LR, Goldstein RZ, Borsook D. The cerebellum and addiction: insights gained from neuroimaging research. Addict Biol. 2014;19(3):317–317.

15. Carta I, Chen CH, Schott AL, Dorizan S, Khodakhah K. Cerebellar modulation of the reward circuitry and social behavior. Science. 2019;363(6424).

16. Abreu-Villaca Y, Seidler FJ, Slotkin TA. Impact of adolescent nicotine exposure on adenylyl cyclase-mediated cell signaling: enzyme induction, neurotransmitter-specific effects, regional selectivities, and the role of withdrawal. Brain Res. 2003;988(1-2):164–172.

17. Chellian R, Behnood-Rod A, Wilson R, et al. Adolescent nicotine and tobacco smoke exposure enhances nicotine self-administration in female rats. Neuropharmacology. 2020;176:108243.

18. Burstein O, Doron R. The Unpredictable Chronic Mild Stress Protocol for Inducing Anhedonia in Mice. J Vis Exp. 2018(140).

19. Nestler EJ. Transcriptional mechanisms of drug addiction. Clin Psychopharmacol Neurosci. 2012;10(3):136–136.

