## Supplementary materials for "AB-Free Kava Suppresses Tobacco Smoke-Induced CREB Phosphorylation in the Mouse Cerebellum"

**Animal Studies Design**

**Round 1** had a 2-week continuous tobacco smoke exposure. Mice in Group 1 were exposed to filtered air and mice in Groups 2 and 3 were exposed to tobacco smoke. Mice in Groups 1 and 2 were maintained on a standard diet while mice in Group 3 were maintained on AB-free kava-supplemented diet (0.75 mg kavalactones/g diet). All mice (air-control diet, n=4; smoke-control diet, n=4; smoke-kava diet, n=4) were euthanized 24h after the last tobacco exposure.

**Round 2** had a 4-week continuous tobacco exposure. Mice in Group 1 were exposed to filtered air and mice in Groups 2 and 3 were exposed to tobacco smoke. Mice in Groups 1 and 2 were maintained on a standard diet while mice in Group 3 were maintained on AB-free kava-supplemented diet (0.75 mg kavalactones/g diet). All mice (air-control diet, n=3; smoke-control diet, n=3; smoke-kava diet, n=3) were euthanized 24h after the last tobacco exposure.

**Round 3** had a 9 (3x3)-week tobacco exposure study with 3 weeks of breaks in between the three-week tobacco smoke exposure. All mice were also given 2 µmol NNK via i.p. injection with tobacco exposure started 24h after. Mice in Group 1 were exposed to filtered air, and mice in Groups 2 and 3 were exposed to tobacco smoke for 3 weeks. Then all mice were housed without any treatment for 3 weeks. The 3-week exposure and 3-week break were repeated three times. Mice in Groups 1 and 2 were maintained on a standard diet while mice in Group 3 were maintained on AB-free kava-supplemented diet (0.25 mg kavalactones/g diet). All mice (air-control diet, n=3; smoke-control diet, n=3; smoke-kava diet, n=3) were euthanized at the end of the 3^rd^ cycle (i.e. 3 weeks after the last tobacco exposure).

**Round 4** had an 11 (3x3 + 2)-week tobacco smoke exposure with breaks in between smoke exposure with intermittent stress paradigm. Mice in Group 1 were given 100 µL saline and mice in Groups 2 and 3 were give 4 µmol NNK via i.p. injection at the start of the experiments. Twenty-four hours after NNK treatment, all mice were exposed to tobacco smoke for 3 weeks. Then mice in Group 1 were housed in the facility without any treatment for 3 weeks, while mice in Groups 2 and 3 underwent stress paradigm treatment (3 times/week, details described in the stress paradigm section) for 3 weeks. The 3-week exposure and 3-week stress (break) were repeated three times, after which all mice were housed in the facility without any treatment for 4 weeks. Mice in groups 2 and 3 were exposed to stress paradigm for 1 week, followed by tobacco smoke exposure for one week, stress paradigm for one more week, and tobacco smoke exposure for one more week. In the meanwhile, mice in Group 1 were exposed to filtered air for 4 weeks. Mice in Groups 1 and 2 were maintained on a standard diet while mice in Group 3 were maintained on AB-free kava-supplemented diet (0.25 mg kavalactones/g diet). All mice (air-control diet, n=4; smoke-control diet, n=4; smoke-kava diet, n=4) were euthanized at 24h after the last tobacco exposure.

| Monday | Wednesday | Friday |
| --- | --- | --- |
| A | B | C |
| D | E | F |
| G | H | I |

A: Physical restraint (15 min)

B: Dampen bedding (2hr) (Dampen the bedding by pouring 10-20 oz. of clean water into each standard clean cage.)

C: Empty cage (Remove bedding from each cage for 2 hr)

D: Forced-swimming test (15 min) (A depth of 30cm is commonly recommenced, although less depth may be adequate for mice.)

E: Tilt cage (Tilt cages to approximately 45° (without bedding) for 2 hr.)

F: Wet cage (Remove all bedding from each cage and add water to a depth of ~0.25 inches for 2 hr)

G: Confined space exposure (0.5 hr)

H: Cage vibration

I: Novel cage exposure (mice were placed on a clean standard rat cage for approximately 1 hr)

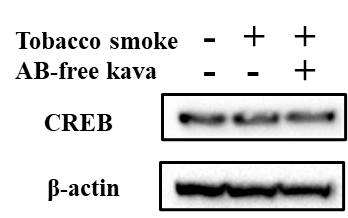

Fig. S1. Representative western blot analysis of cerebellum total CREB upon TS exposure with/without AB-free kava supplementation (n=3).
